# Perspective matters: using past experiences to understand others’ insensitivity to physical pain

**DOI:** 10.1101/2025.07.17.665290

**Authors:** Federica Meconi, Marco Canducci, Sebastian Michelmann, Alessandro Grecucci, Irene Sperandio

## Abstract

Prior research has shown that autobiographical memory (AM) supports empathy, allowing individuals to use the self as a model for understanding others. However, when the subjective experience of an event diverges between observer and target, relying on AM may hinder empathic processes, rather than support them.

In this study, we recorded electroencephalographic (EEG) data from 36 healthy young adults during a pain decision task to investigate empathy for individuals described as either sensitive to physical pain (like the participants) or clinically insensitive to physical pain. Participants reported lower empathy for insensitive than for sensitive targets, regardless of whether they had personally experienced the specific event that caused physical pain.

Event-related potential (ERP) analyses revealed differences in neural responses to sensitive and insensitive targets at a late stage of processing, in the discending phase of the P300 component; these differences were positively correlated with individual differences in empathy.

To determine the underlying mechanisms, we applied multivariate pattern analysis (MVPA) to assess whether empathy for physical painrelies on AM reactivation or instead engages active perspective taking.

We observed that AM reactivation can serve as a flexible mechanism that supports empathy by enabling observers to shift perspective when direct experiential overlap is lacking.

## Introduction

Mentalizing, the ability to attribute mental states to oneself and others, is central to human social cognition. Effective interpersonal understanding depends on the ability to adopt another person’s perspective (perspective taking) or to internally simulate their emotional state (empathy) while maintaining a clear distinction between self and other (Quesque et al., 2024). Several meta-analysis have confirmed that partially dissociable anatomical networks subserve these processes (Lamm et al., 2011; Molenberghs et al., 2012, 2016). Event-related potentials (ERPs) studies further supported this distinction, revealing different temporal dynamics for empathy and perspective taking (Fan and Han, 2008; Sessa et al., 2014a, 2014b; Sessa and Meconi, 2015; Meconi et al., 2018; Palmieri et al., 2021).

Recent research suggests that autobiographical memory (AM), plays a crucial role in supporting mentalizing processes (Spreng and Grady, 2009; Meconi et al., 2021). By leveraging stored personal experiences, individuals can use the self as a model for inferring others’ internal states. However, this strategy may not be universally adaptive: the observer’s subjective experience may diverge from the target’s if the emotion associated with a past experience does not align with the target’s current emotional state.

Evidence from clinical populations underscores this complexity. For instance, Danziger and colleagues (Danziger et al., 2009) demonstrated that individuals with congenital insensitivity to pain exhibit brain activation patterns of the anterior cingulate cortex comparable to that of healthy controls when observing others in physical pain. This suggests that autobiographical experience of physical pain is not necessary for an empathic neural response. However, these individuals reported lower subjective rates of empathy, revealing a dissonance between self-reported experience and the neural markers of empathy.

Building on this premise, the present study combines electroencephalography (EEG) with machine learning techniques (Multivariate Pattern Analysis, MVPA) to investigate whether AM with an associated emotion that is incongruent with the target’s experience is reactivated despite it being uninformative to infer others’ mental states, or it is rather flexibly inhibited to support accurate perspective taking. We used events of physical pain that were confirmed to belong to participants’ AM and that could not be internally simulated or imagined if not part of the observer’s past experiences, i.e. the fracture of a bone.

Recent advances in MVPA have demonstrated that brain activity patterns can be tracked during the encoding of new neutral episodes and re-activated during their retrieval (Michelmann et al., 2016; Linde-Domingo et al., 2019). Also, in a recent study Meconi et al. (2021), using a similar MVPA approach, provided the first direct evidence for the online reactivation of AM when participants were required to explicitly rate their empathy for others’ pain. In that study, participants initially performed a pain decision task in which they were asked to emapthize with painful or neutral experiences for which they did or did not have an associated AM. In a subsequent task, shapes were used to prompt a mind representation of the scenes described during the empathy task. A cross-task classification showed evidence of an AM reactivation when empathizing with others’ physical pain. In the current study, we recorded EEG while participants performed a similar paradigm. A first pain decision task was used to elicit an empathic reaction for targets identified as being sensitive (healthy like the participants) or clinically insensitive to physical pain. Targets were preceded by sentences describing events that cause physical pain that participants had either experienced (AM) or not (non-AM). In the second task, participants were asked to actively visualise in their mind’s eye all the physical pain events showed in the pain decision task from either a first-person or third-person perspective. This retrieval task, henceforth classifier task, was designed to extract the neural fingerprints of AMs and non-AMs as well as those associated with the first- and third-person perspectives. A linear discriminant analysis (LDA) classification (Fisher, 1936) was trained on the EEG data from the classifier task and tested on data obtained from the pain decision task. We sought to test the spontaneous reactivation of AMs or the ability to take the other’s perspective when participants were asked to empathize with targets, either sensitive or insensitive to physical pain, described in events that participants did or did not experience themselves in the past.

In doing so, we observed the EEG pattern associated with perspective switching from the first-to the third-person when participants empathized with others’ physical pain.

## Methods

The study was approved by the University of Trento Research Ethics Committee (2021-034). Written consent was obtained from all participants. All the participants were recruited through social media web pages of the University of Trento and obtained a cash reimbursement for their time (€10/h). They had normal or corrected-to normal vision. The inclusion criteria entailed excellent Italian proficiency, having experienced events that caused intense physical pain, no history of neurological or psychiatric disorder, and no use of psychoactive drugs for medical or recreational purposes.

### Participants

We collected data from forty-two participants (9 males, mean age 22 y/o, sd = 3, 3 left-handed). We calculated a priori minimum sample size of 36 participants, based on G*Power (Faul et al., 2007) calculations to attain a moderate effect size with α = 0.05 and power = 0.95 for a 2×2 repeated measures ANOVA. Six participants were discarded from the final sample because they did not complete the tasks. Therefore, the final sample consisted of 36 full datasets (5 males, mean age=21.8 y/o, SD=2.7, 3 left-handed).

### Questionnaires

Before the experimental session, students who expressed interest in participating in the experiment completed a questionnaire to ascertain whether or not they had experienced events of intense physical pain in the past. We compiled a list of 32 events and asked participants to indicate if the event had occurred, how many times it had occurred, and to rate the intensity of the pain they perceived if it had occurred, or the intensity they believed they would experience if such an event were to happen to them. We aimed to collect eight events per participant with comparable pain intensity: four for which participants had an associated AM and four for which they did not. Once elegibility was determined, participants were invited to an experimental session in the lab and sent a link to complete a series of questionnaires. The questionnaires were administered in a randomized order in Qualtrics and assessed participants’ dispositional empathy resources (using the Empathy Quotient, EQ; Baron-Cohen and Wheelwright, 2004); the Interpersonal Reactivity Index, IRI; Davis, 1983;) as well as their ability to recognize and describe their own emotions (using the Toronto Alexithymia scale, TAS-20; Bagby et al., 1994). On average, participants scored within the normal range of the EQ (M = 48.94 SD = 9.55), showed no evidence of alexythimia as indicated by the TAS score (M = 55.28, SD = 6.52), and did not exhibit traits associated with the autism spectrum in the general population (M = 16.44, SD = 5.44). The IRI scores are reported in Table 1.

**Table 1.** Mean scores and standard deviation (SD) of the Interpersonal Reactivity Index (IRI)’s subscales.

|  | *Mean* | *SD* |
| --- | --- | --- |
| **FS** | 26 | 5,03 |
| **PT** | 26,81 | 3,87 |
| **EC** | 27,36 | 4,09 |
| **PD** | 20,11 | 4,73 |

### Stimuli and Procedure

All the stimuli were presented on a grey background (RGB: 122 122 122) of a 24’‘computer screen with a refreshing rate of 70 Hz and a display resolution of 1980×1080 pixels. The experimental tasks were programmed using Psychtoolbox-3 (Kleiner, 2010) in Matlab.

Participants sat comfortably on a chair at 70 cm from the computer screen in a dim light room. They wore an elastic cap with EEG electrodes while performing two sequential tasks that lasted around 80 minutes including the practice trials. For all participants, the first task was the pain decision task, followed by a classifier task.

#### The pain decision task

##### Stimuli

Stimuli consisted of sentences, describing events that involved intense physical pain, each followed by faces (see Fig. 1A). The faces comprised 16 identities with neutral facial expression, 8 males and 8 females. The faces were presented in greyscale and were equalized for luminance using the SHINE toolbox (Willenbockel et al., 2010). Each face was surrounded by a colored frame, either blue or yellow with RGB [255 240 0] or [0 0 255] respectively, counterbalanced across participants, to indicate whether the target of empathy was sensitive to pain or suffered from a rare clinical condition known as Congenital Insensitivity to Pain. The sentences were written in black Helvetica font, size 20, and were tailored for each participant so that four of the events were associated with an AM, while the others were not. All sentences followed the same structure, either “This person got – […]” or “This person did – […]”, to ensure consistent syntactic complexity.

**Fig 1.**
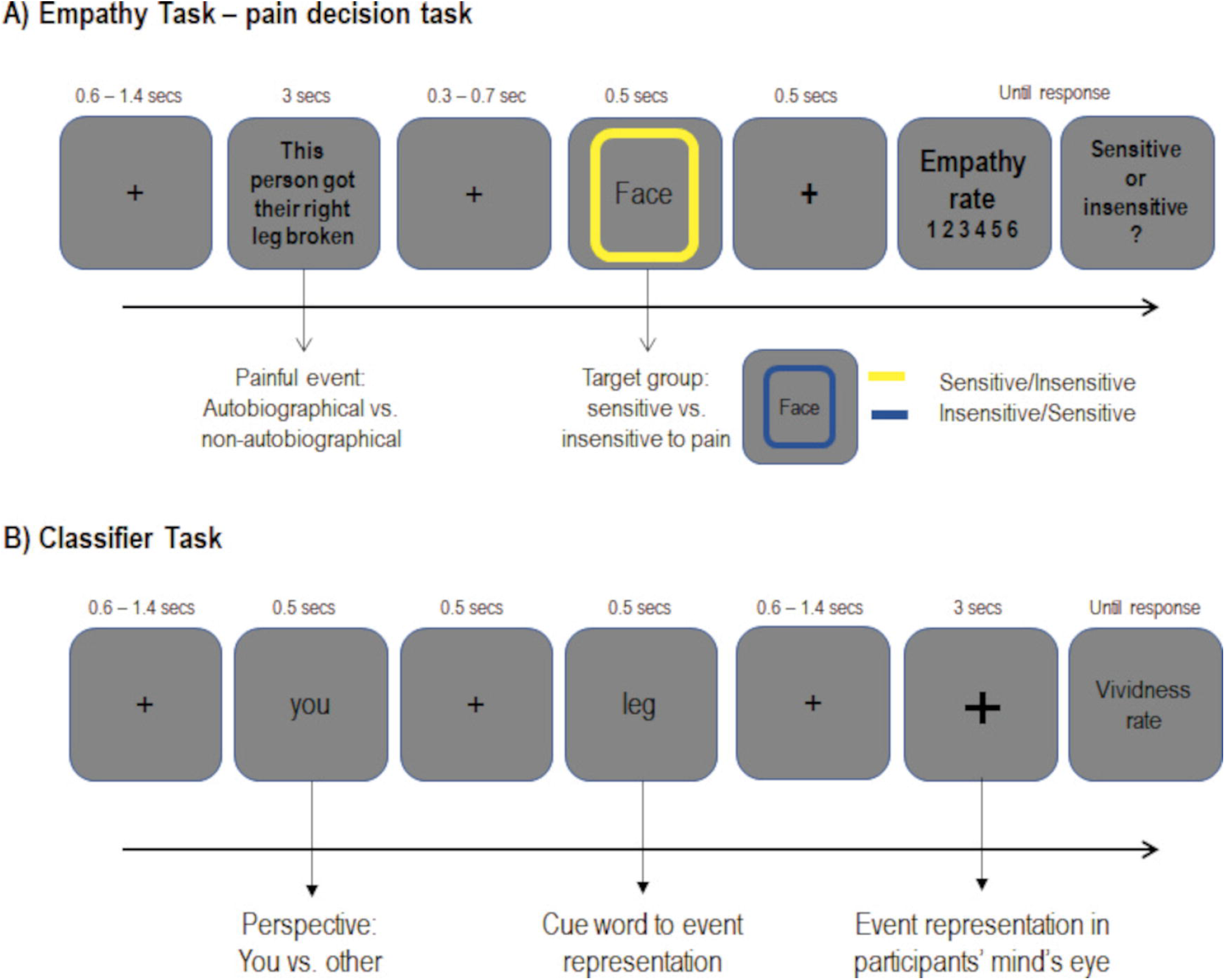
Schematic representation of the experimental paradigm. **A)** The trial scheme adopted in the empathy inducing pain decision task. Blue and yellow frames surrounding the neutral faces were counterbalanced across participants to indicate sensitive or insensitive targets. **B)** The trial scheme adopted in the cliassifier task.

##### Paradigm

Each trial started with a fixation cross displayed for a variable duration (600-1400ms in 100ms steps). Participants were then presented with the sentence (3s) followed by a face (500ms) interleaved by a variable fixation cross (800-1600ms, jittered in 100ms steps). For each trial, participants were asked to rate their empathy for the person just presented and described within the specified preceding context on a scale ranging from 1 (“Not empathic at all”) to 6 (“Very much empathic”). The rating was self-paced and made by pressing one of six response keys on the computer keyboard: “s”, “d”, “f”, “j”, “k”, “l”, with “s” corresponding to “1” and “l” to “6”. The rating was provided after another fixation cross displayed for 500ms. Finally, participants had to indicate whether the target was sensitive or insensitive to physical pain by pressing “j” or “f” on the keyboard and only correct trials were analysed. The button assignments for indicating sensitivity or insensitivity to pain were counterbalanced across participants, and the correspondence between the key and the answer was always shown on the screen. The task consisted of 48 trials per condition: AM vs. non-AM contexts followed by sensitive vs. insensitive targets. Trials were pseudo-randomized to ensure that conditions were balanced over the full session. There were 192 trials in total divided into 4 blocks preceded by 6 practice trials. A schematic representation of a trial is depicted in Fig.1A.

#### The classifier task

##### Stimuli and Paradigm

Participants were asked to visualize in their mind’s eye one of the events previously presented in the pain decision task, using either a first- or third-person perspective. Each trial started with a variable fixation cross (600-400ms, jittered in 100ms steps). Participants were then presented with the word “You” or “Other” (500ms), which cued them to imagine the event either from a protagonist’s (first-person) or observer’s perspective (third-person). Next, a cue-word (500ms) was presented after a second fixation cross (500ms) to indicate which of the eight possible events they had to visualize. For example, we used “arm” as a cue-word for the sentence “This person got their right arm broken”; “ligament” for “This person got their ligament torn” and so on. Text was presented in black Helvetica, size 20. After a third variable fixation cross (600-1400ms jittered in 100ms steps), a thicker black fixation cross was displayed for 3s to cue participants to start visualizing the event. Finally, participants were asked to rate the vividness of their visualization on a 1-6 points scale, with “1 = Not vivid at all” to “6 = Very vivid”. The rate was self-paced and made by pressing one of six response keys on the computer keyboard: “s”, “d”, “f”, “j”, “k”, “l”, with “s” corresponding to “1” and “l” to “6”. This task served as a localizer for an EEG pattern classifier to extract the neural fingerprints of both the memory scenes and the perspectives, which were later used to probe data from the pain decision task.

The task was preceded by 8 practice trials. Participants were allowed to repeat the practice block until the task and the association between the cue-words and the related scenes was clear to the participants. There were 60 trials per condition: AM vs. non-AM scenes represented from a first- and third-person perspective, that were pseudo-randomized to ensure a balanced distribution across the 240 trials comprising the full session. The classifier task, depicted in Fig.1B, was divided into 4 blocks.

### Data acquisition and analysis

The EEG was recorded using a BrainProducts EasyCap from 64 Ag/AgCl passive electrodes placed according to the international 10-10 system. EEG signal was recorded with a band-pass filter of 0.05 – 100 Hz and digitized at a sampling rate of 1024 Hz using BrainVision Recorder software (BrainProducts). Impedance was kept below 10 kΩ and regularly checked during the experiment. The ground electrode was placed on AFz. The online reference was placed on FCz and the EEG was then re-referenced offline to the average reference. EEG data was segmented into epochs of 3s, starting 1s before the onset of the face for the pain decision task and of 6s, starting 2s before the cue onset for the classifier task. The epoched data was visually inspected to discard large artifacts from further analysis. Further preprocessing steps included Independent Component Analysis for ocular artifacts correction and average interpolation of noisy channels when needed. After removing trials contaminated by eye and muscle artefacts a comparable number of trials remained for all conditions (see Supplementary Materials for further details). EEG data was analyzed with MATLAB (©Mathworks, Munich, Germany) using the open-source FieldTrip toolbox (Oostenveld et al., 2011) and in-house Matlab routines.

### Univariate ERP analysis

#### Pain Decision task

Cluster-based permutation tests were performed over the whole scalp and within a 1s time-window on ERPs time-locked to face onset. This analysis aimed to account for any significant differences between AM and non-AM contexts, as well as between sensitive and insensitive targets to reveal ERP empathic responses to the contexts or faces. Additionally, a preliminary analysis was carried out to test for any involvement of memory in the empathy task. To this end, cluster-based permutation tests were performed on the ERPs time-locked to the onset of the sentence contrasting AM vs. non-AM trials.

### Linear Discriminant Analysis EEG pattern classifier

#### Classifier task

Linear Discriminant Analysis (LDA) is a MVPA method that identifies a decision boundary to distinguish patterns of brain activity associated with different stimulus categories. Decision values are an index of the distance of each classified category from the hyperplane that represents the decision boundary. This method is based on specified features of the EEG signal (e.g., amplitude, frequency etc.). LDA can then estimate, with a certain accuracy, whether the pattern of brain activity in new data, not used during the training of the classifier, matches more closely one stimulus category or the other.

In the current study, the LDA was trained and tested on the EEG raw patterns (i.e. amplitude of the signal on each electrode), for each participant at each time point, and regularized with shrinkage (Blankertz et al., 2011). We equalized the number of trials for AM and non-AM events and for first and third-person perspectives before training the classifier to ensure that the output was not biased by any slight difference in the the signal-to-noise ratio.

In order to reduce unwanted noise and computational time, the signal was filtered between 0.1 and 40Hz and down sampled to 128Hz before classification with a baseline correction window of 500ms before the onset of the event visualization (the “Event representation in participants’ mind eye” in Fig.1B).

To ensure that the task could successfully distinguish between the EEG pattern reflecting AM and non-AM as well as between the first- and the third-person perspective, we ran a sanity check of the LDA on the classifier task. The classifier was trained and tested with a K-fold cross-validation procedure with K = 5 during the event visualization. For each participant, a set of two (generalization) matrices was extracted, containing each the decision values for the given condition, once to distinguish between memory scenes and once to distinguish between perspectives. For each participant, we tested whether the diagonal elements of these matrices represented different conditions.

Once determined that the task successfully distinguished the EEG patterns in all conditions, we tested it during the empathy task. In this second step, the classifier was trained during the event visualization in the classifier task and then tested on the empathy task in a 1s time-window starting from the face onset. Crucially, any consistency in neural patterns observed across tasks would indicate the representation of the memories or perspectives during the empathy task.

The aim of this cross-classification was to test whether participants’ brain activity differentiated between AM and non-AM scenes, or whether they switched perspective between the first- and the third-person when empathizing with others’ insensitivity to physical pain.

### Statistical analysis

Mean proportions of the empathy rating scores of correct trials (i.e., trials in which participants correctly identified the target as sensitive or insensitive to pain) were computed for each condition and analyzed in a 2×2 repeated measures ANOVA with Memory (Autobiographical vs. non-Autobiographical) and Target (Sensitive vs. Insensitive) as within-subjects factors for the pain decision task. For the classifier task, a 2×2 repeated measures ANOVA with Memory and Perspective (First vs. Third) as within-subjects factors was carried out on vividness scores. Tukey corrected paired-sample t-tests were conducted when appropriate to further explore significant interactions.

For the sanity check of the classifier we extracted the diagonals of each participant’s set of two generalization matrices containing each the decision values for the memory and perspective conditions. The diagonals of these matrices are the decision values of the classifier trained and tested at the same timepoint and are therefore constituted of timepoints that are independent from each other. To test whether the distributions of diagonal elements for each pair of conditions were significantly separable for each participant, we carried out a t-test on these samples.

For the across-tasks classification, an empirical null distribution was created using a combined permutation and bootstrapping approach (Stelzer et al., 2013) to test whether the maximum cluster of accuracy values above the chance level was statistically significant. Clusters were identified on the basis of the number of adjacent pixels found with the Matlab function bnlabel. We used the LDA in 100 matrices with pseudo-randomly shuffled labels independently for each participant and created a null distribution of accuracy values that we contrasted with the LDA outputs obtained with the real data. This was done by sampling with replacement 100000 times from the real and random data of each participant and computing a group average. This procedure resulted in an empirical chance distribution, which allowed us to investigate whether the results from the real-labels classification had a low probability of being obtained due to chance (p<0.05) (i.e., exceeding the 95^th^ percentile).

## Results

### Behavioural results – Empathy task

The ANOVA revealed a main effect of Target on empathy judgements, with higher ratings for sensitive than insensitive targets (*F*(1,35)=54.441; *p*=.000000012, *η*_*p*_^2^ = .609). The main effect of Memory was not significant (*F*(1,35) = 1.590; *p* = .216, *η*_*p*_^2^ = 043) although slightly higher ratings for AM than non-AM scenes. The interaction between Memory and Target did not reach significance (*F*(1,35)=2.586; *p*=.117, *η*_*p*_^2^=.069). Results are represented in Fig 2A.

**Fig 2.**
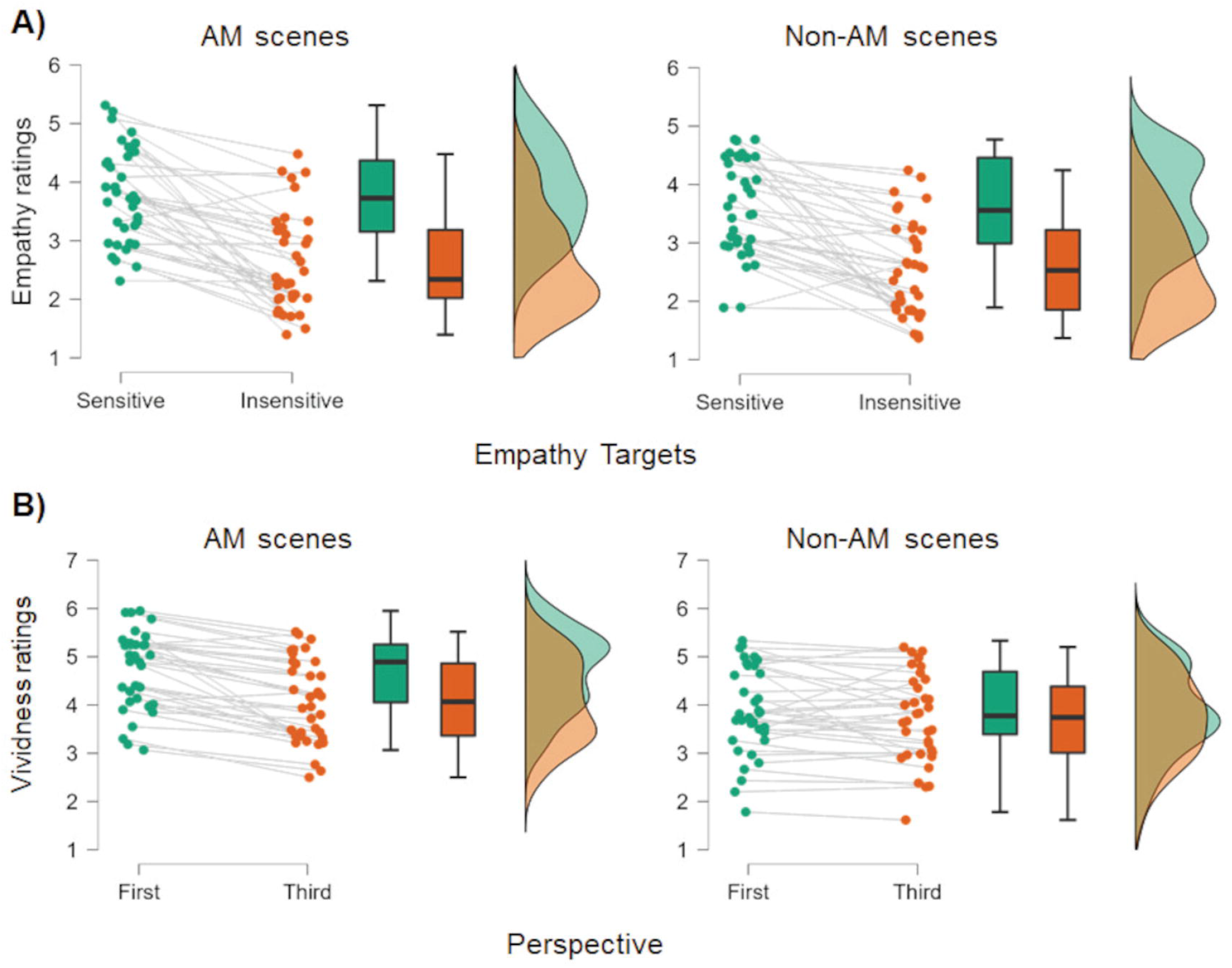
Behavioral results. **A)** Results from the pain decision task carried out on the empathy rates. **B)** Results from the classifier task carried out on the vividness rates.

### Behavioural results – Classifier task

The ANOVA revealed a main effect of Memory (*F*(1,35)=27.529; *p*=.0000076, *η*_*p*_^2^=.440) as well as a main effect of Perspective (*F*(1,35)=26.268; *p*=.000011, η_*p*_^2^=.429) on vividness scores. Participants mentally represented AM scenes more vividly than non-AM scenes. They also reported greater vividness for scenes imagined from a first-compared to a third-person perspective. The interaction between Memory and Perspective was also significant (*F*(1,35)=29.031; *p*=.000049, *η*_*p*_^2^=.453). Post-hoc comparisons revealed that participants represented AM scenes more vividly from a first-compared to a third-person perspective (*t*(35)=7.114, *p*_*c*_=.000000014), whereas vividness did not differ between perspectives for non-AM scenes (*t*(35)=1.809, *p*_*c*_=.28). Results are presented in Fig 2B.

### EEG results

#### ERPs

Cluster analysis conducted over a 1s time-window, from the onset of the face until the presentation of the rating scale, revealed one right posterior cluster of electrodes showing a significant ERP difference between sensitive and insensitive targets between 550-750ms (*p*_c_=0.022). Fig.3A depicts the ERPs showing the significant difference between sensitive and insensitive targets, and Fig. 3B the topography of its significant cluster. Cluster analysis conducted over a 3sec time-window, during the preceding sentence presentation, did not reveal any significant cluster (*p*_c_>.05).

**Fig 3.**
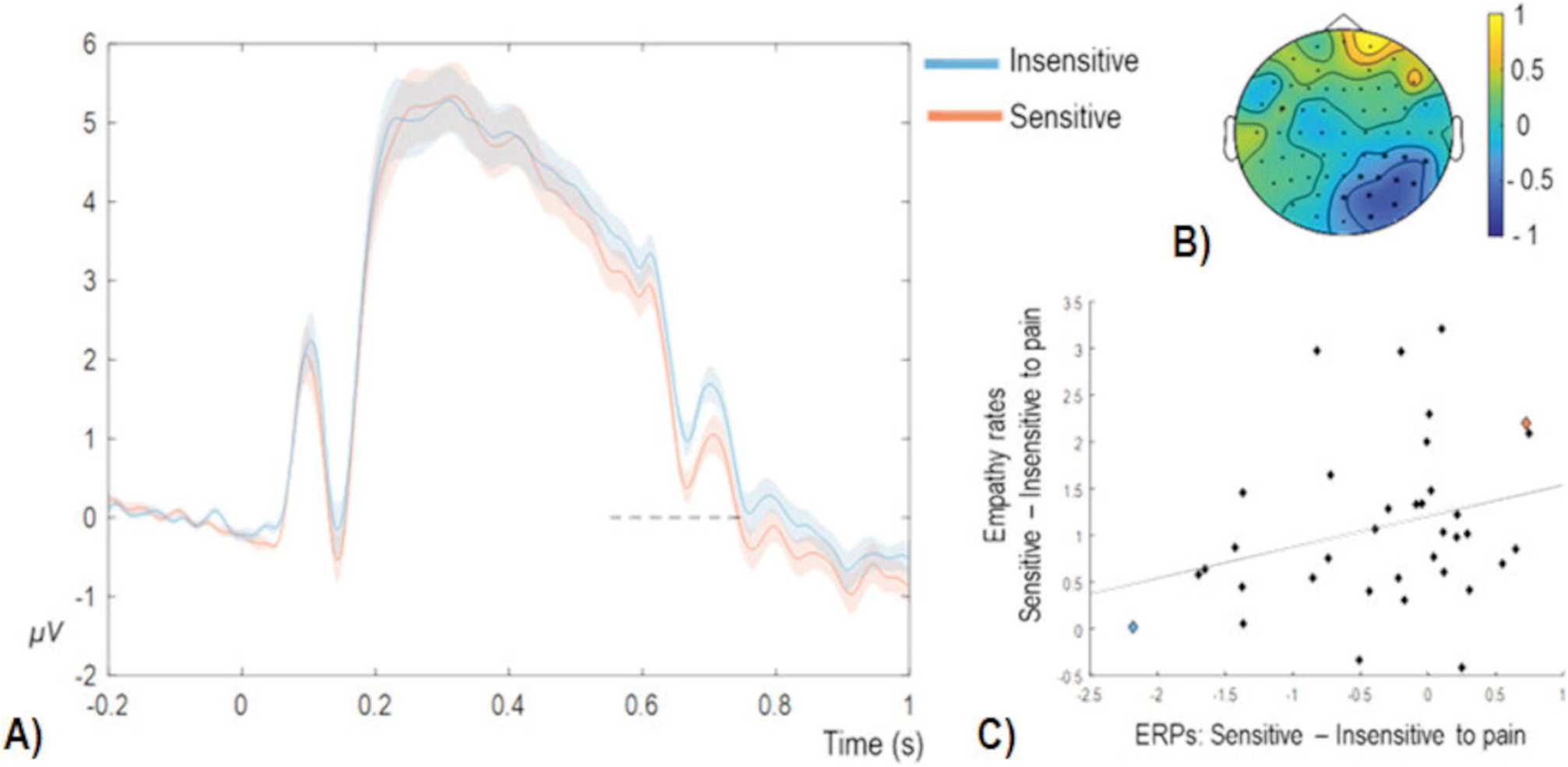
ERP findings. **A**) ERPs time-locked to the onset of the face and reflecting sensitive and insensitive to physical pain targets of empathy at the posterior. **B)** Clusters analysis performed over all the electrodes in a 0 – 1 sec time-window. Colors code t-values. **C)** Correlation between differential scores of the subjective rates of empathy, y axis, and the differential scores of the ERPs, x axis.

To examine a possible relationship between ERPs and subjective ratings of empathy, ERPs and ratings for insensitive targets were subtracted from those for sensitive targets. We observed a positive correlation, *r*=.28, *p*=.05, between these differential scores in a pool of posterior electrodes, including the significant cluster, during a sustained time-window (600-800ms) shown in Fig 3C. Positive values for both indices indicated greater empathy for sensitive compared to insensitive targets. Therefore, this correlation suggests that the greater the ERP amplitude for sensitive compared to insensitive targets, the higher participants rated their empathy for sensitive than insensitive targets.

#### LDA

The sanity check for the classifier task showed that decision values differed significantly for 30 out of 36 participants (max. *p*=.012) when classifying mental representations of AM versus non-AM events. Similarly, decision values differed significantly for 32 out of 36 participants (max. *p*=.03) when classifying first-versus third-person perspectives. The distributions of decision values for the memory and the perspective conditions for each participant are shown in the Supplementary Materials: Figs. S1 and S2, respectively. The bootstrapping analysis carried out for the across-tasks classification revealed a significant cluster (*p*=.0014 with all 36 participants; *p*=.0019 without the four non-classifying participants) within a sustained time window (0.25-1secs), providing evidence for the online switching of perspective. In contrast, no significant cluster was found for memory reactivation (*p*=.94 with all 36 participants; *p*=.96 without the six non-classifying participants), suggesting that memory content was not reactivated in preparation for the empathy judgement. The classifier achieved a peak accuracy of 0.53 for the significant cluster, which was meaningful although modest. Fig. 4 shows the cluster of accuracy obtained from the across-tasks classification for all participants that was significantly above the chance level.

**Fig 4.**
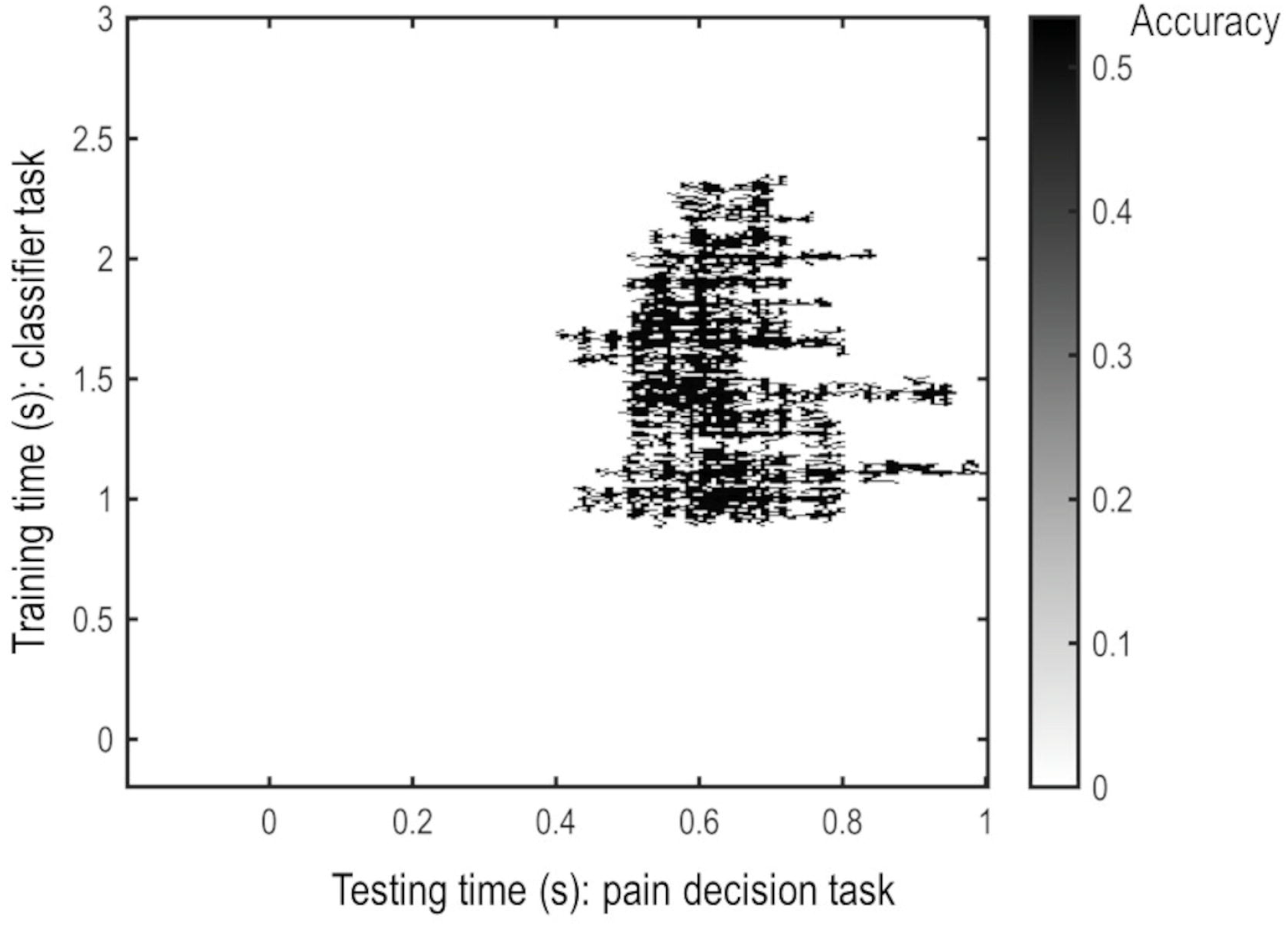
LDA results. Time by time generalization matrix obtained by training the LDA during the classifier task (timing represented on the y axis) and tested during the pain decision task (timing represented on the x axis). Colorbar shows the accuracy of the LDA. Cluster only shows the significant accuracy above chance level.

## Discussion

This study investigated the role of AM in empathy when observers and targets shared the same physical pain experience but associated it with divergent emotional states. EEG was recorded during both an empathy task and a subsequent classifier task in which participants had to represent in their mind’s eye the AM and non-AM scenes from a first- or third-person perspective. An LDA classifier was trained on EEG data from the classifier task and applied to the empathy task to determine whether empathy would engage AM reactivation or perspective switching when one’s own past experiences would be uninformative to understand others’ emotional state. Our findings provided evidence for active perspective switching when healthy participants evaluated the pain of others described as sensitive or clinically insensitive to physical pain.

In the pain decision task, participants reported higher empathy for sensitive, compared to insensitive, targets and ERPs mirrored this finding by showing a significant differentiation between targets at a late stage of processing, around 550-750ms after face onset. Previous ERP studies on empathy for pain have generally shown that an empathic reaction is expressed as a positive shift in the ERP response for painful compared to a neutral condition (e.g., Meconi, Doro, Lomoriello, Mastrella, & Sessa, 2018; Sessa & Meconi, 2015). We observed a positive shift in the ERPs for insensitive, compared to sensitive, targets within a 1s time-window from a cluster of posterior electrodes. However, the differential mean amplitudes of this response positively correlated with the differential scores of participants’ subjective ratings of empathy. Therefore, the group ERP results suggest that this difference was associated with an increased level of empathy for physical pain experienced by individuals who could perceive physical pain whereas this resonance with the others’ pain was appropriately reduced for those described as insensitive to pain. This correlation underscored a close alignment between self-reported empathy and its underlying neural processing.

In the classifier task, behavioral results showed that participants represented the events more vividly from a first-than a third-person perspective, particularly for the AM scenes. Noteworthy, decoding the imagined perspective across tasks could have been particularly challenging since participants were never explicitly prompted to switch perspective during the empathy task and no perceptual cue aided the mind representation of the perspective nor of the events. Stecher and Kaiser (2024) recently used an LDA classifier across tasks and showed the feasibility of decoding imaginary elements of a scene even in the absence of direct visual presentation. Crucially, MVPA revealed that participants engaged in active perspective switching rather than automatically reactivating their own memories when direct experiential overlap was lacking between observer and target. This finding suggests that participants were able to inhibit reliance on AM when such retrieval would interfere with accurate others’ understanding.

Several methodological features further reinforce the robustness of our findings. First, the experimental design deliberately avoided emphasizing the autobiographical component of the scenes as well as the active perspective switching. This allowed us to assess whether AM would be spontaneously reactivated in the absence of explicit prompting or whether participants would take the others’ perspective to attenuate their empathic response towards the physical pain undergone by someone who is insensitive to pain. Second, the classifier task was administered always after the empathy task to minimize the possibility that participants would intentionally either recall personal memories or activate perspective taking processes during the empathy task. Third, we controlled for perceptual confounds by using different types of stimuli for the two tasks: empathy was elicited with sentences and faces, while memory representations were cued by fixation crosses, preceded by cue-words. Importantly, the same structure was applied across all four conditions, ensuring that classifier outputs reflected cognitive processes (e.g., memory or perspective switching) rather than perceptual features. It is important to notice that the accuracy peak of the classifier was modest. However, we identified a significant cluster indicating that consistent information was shared across tasks. Critically, this shared EEG pattern cannot be attributed to the perceptual features of the stimuli used for training and testing the classifier because of the methodological features of our design. These findings align with previous studies that have successfully decoded neural patterns during a stage of processing where information was represented mentally in absence of any perceptual feature (Kurth-Nelson et al., 2015; Hebart and Baker, 2018; Kerrén et al., 2018).

In contrast with previous findings that have shown that past experiences support empathic abilities (Perry et al., 2011; Bluck et al., 2013; Gaesser and Schacter, 2014; Meconi et al., 2021), we found no evidence that AM reactivation guided empathy processes. While these findings may initially appear divergent with previous studies, they collectively suggest that if shared experiences can potentially inform empathy, individuals are capable of modulating their perspective-taking strategies. This modulation allows them to prioritize the emotional state of the target over their own autobiographical associations, a flexibility that may be crucial for achieving accurate and context-sensitive social understanding.

Furthermore, this picture can contribute to explain the puzzling findings that neuropsychological studies report concerning empathy resources following memory loss (Meconi et al., 2021b). In fact, if memory is necessary to empathise then clinical conditions associated with memory deficits would consistently lead to evidence for empathy impoverishment. Studies in patients with degenerative disorders that involve memory loss (e.g., Monetta et al., 2009; Duval et al., 2012; Moreau et al., 2013, Moreau et al., 2015; Pell et al., 2014) support a co-occurrence of memory and empathy deficits. However, these conditions often involve a global cognitive decline, which cannot clarify the causal relationship between these deficits. The handful of studies that examined different empathic abilities, such as perspective taking or empathy and self-other distinction (Quesque et al., 2024), in clinical populations with focal memory loss, such as hippocampal amnesia, showed spared or only mildly impaired some, but not all, empathic resources (Rosenbaum et al., 2007; Rabin et al., 2012; Beadle et al., 2013; Staniloiu et al., 2013; Sawczak et al., 2019; Demichelis et al., 2020). Healthy aging seems to be associated with progressive empathy decreseas but spared perspective taking (Duval et al., 2011; Chen et al., 2014; Ze et al., 2014). Therefore, empathic processes might appear reduced with progressive AM degeneration, however spared empathic abilities suggest that empathy can leverage other memory resources, in line with studies highlighting the role of acquired semantic knowledge (Mitchell et al., 2005; Pehrs et al., 2017, 2018).

In this vein, research on empathy for pain has consistently shown that affective and semantic associations stored in long-term memory, such as prejudice and stereotypes towards other ethnicities, can shape empathic responses (Xu et al., 2009; Avenanti et al., 2010; Sessa et al., 2014a). Furthermore, rapid formation of new associations, such as pairing a human name with non-human entities, can modulate empathic responses within a short time-window of exposure to the humanized target, i.e. the target paired with a human name (Vaes et al., 2016). These findings provide converging evidence that empathic processing can also be influenced by long-term memories that are not autobiographical in nature (Meconi et al., 2015).

The current study nicely dovetails with this multifaceted scenario in which AM reactivation in empathy is a flexible mechanism. Our findings provide compelling evidence, for the first time, that participants can distance themselves from their own past experiences to actively take the others’ perspectives and understand their emotional state, despite their own subjective experience.

Future works should examine empathy as the result of an interplay between perspective taking and the active inhibition of the autobiographical emotions and its projection onto others.

## Limitations

There are a few limitations that should be mentioned for the current study.

Our sample of participants was mainly composed of female young adults. Although, we did not have any hypothesis about sex differences for our research question, future studies might need to further explore potential differences, either between biological sexes or gender identities.

The accuracy peak we obtained from the across-tasks classification is just above chance. Although modest, the classifier output was meaningful. However, future investigations may adopt a task that could further challenge the LDA or use different classifier methods to increase accuracy.

Finally, the paradigm specifically focused on physical pain. Although physical pain is often employed as a model to study empathy, future studies may expand the investigation to other types of pain or emotions.

## Acknowledgements

This project was funded by the ESRC (NºES/S001964/1) awarded to F.M. M.C is supported by the EPSRC Prosperity Partnerships grant ARCANE, EP/X025454/1. The Authors thank all the participants and the students who helped with data collection.

## Authors contribution

F.M.: conceptualization, data curation, formal analysis, investigation, methodology, funding acquisition, project administration, writing - original draft, writing - review and editing; C.M.: formal analysis, writing - review and editing; S.M.: formal analysis, writing - review and editing; A.G.: methodology, writing - review and editing; I.S.: methodology, writing - review and editing. All authors gave final approval for publication and agreed to be held accountable for the work performed therein.

## Data Availability Statement

A preregistration of the study is available at https://osf.io/qyp48/?view_only=c70fe36ae4124652a9f71c0ca5c04086. The data underlying this article will be shared on reasonable request to the corresponding author.

