## Supplementary Materials for "Perspective matters: using past experiences to understand others’ insensitivity to physical pain"

*EEG analysis.*

Pain Decision Task. After removing trials that were contaminated by eye and muscle artefacts, an average of 89 trials (range: 75 - 96) remained for AM condition, 89 trials (range: 76 - 96) for non-AM condition, 89 trials (range: 76 - 96) for sensitive targets, and 89 trials (range: 76 - 96) for insensitive targets.

Classifier Task. After removing trials which were contaminated by eye and muscle artefacts, an average of 106 trials remained for AM (range: 83-119) and non-AM (range: 81-119) conditions. On average, the same number of trials remained also for first-person (range: 82-119) and third-person (range: 82-119) perspectives.

*Classifier sanity check.*

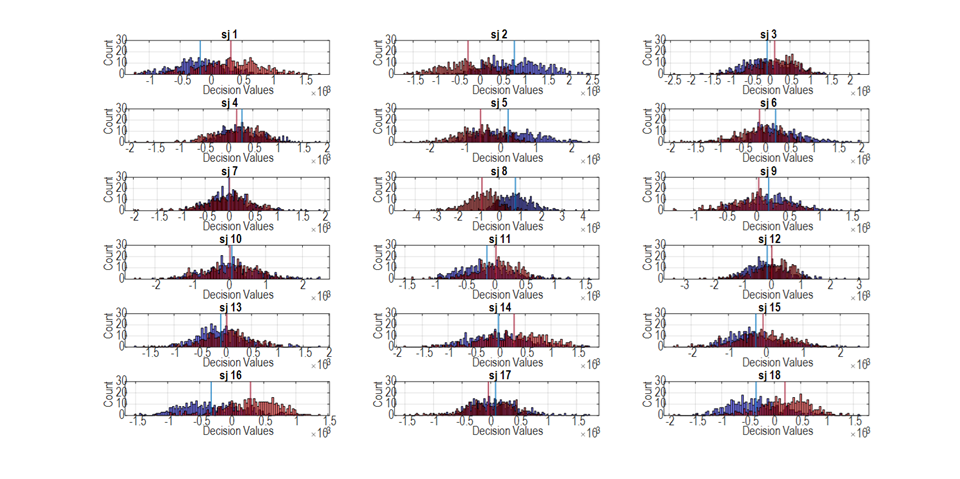

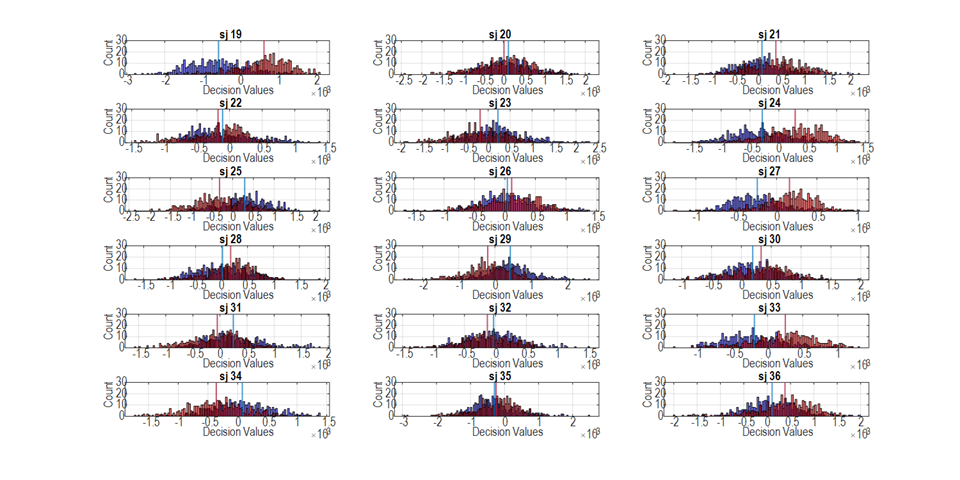

**Fig S1.** Distribution of the decision values extracted from the diagonals of the time by time generalization matrices classifying AM and non-AM scenes retrieval.

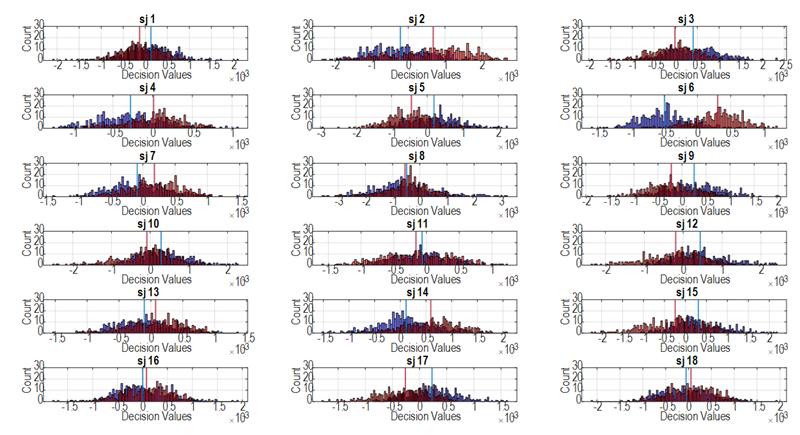

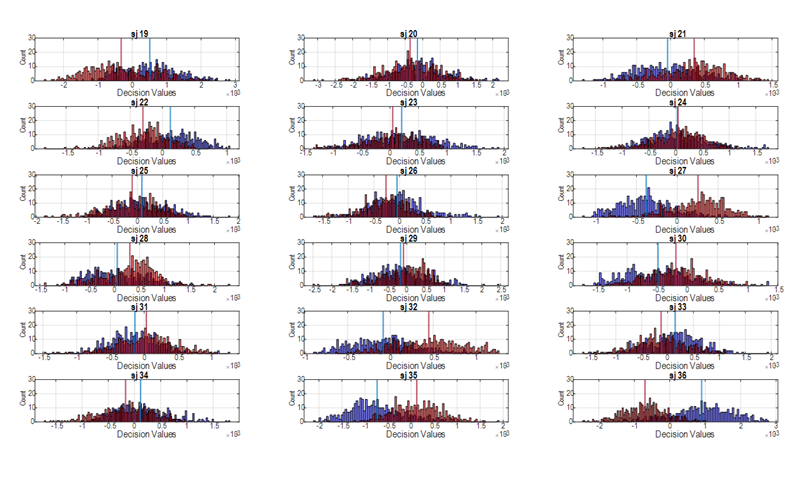

**Fig S2.** Distribution of the decision values extracted from the diagonals of the time by time generalization matrices classifying first- and third-person perspectives.
